# *In-silico* definition of the *Drosophila melanogaster* matrisome

**DOI:** 10.1101/722868

**Authors:** Martin N. Davis, Sally Horne-Badovinac, Alexandra Naba

## Abstract

The extracellular matrix (ECM) is an assembly of hundreds of proteins that structurally supports the cells it surrounds and biochemically regulates their functions. *Drosophila* has emerged as a powerful model organism to study fundamental mechanisms underlying ECM protein secretion, ECM assembly, and ECM roles in pathophysiological processes. However, as of today, we do not possess a well-defined list of the components forming the ECM of this organism. We previously reported the development of computational pipelines to define the matrisome - the ensemble of genes encoding ECM and ECM-associated proteins - of humans, mice, zebrafish and *C. elegans*. Using a similar approach, we report here that the *Drosophila* matrisome is composed of 641 genes. We further classify these genes into different structural and functional categories, including an expanded way to classify genes encoding proteins forming apical ECMs. We illustrate how having a comprehensive list of *Drosophila* matrisome proteins can be used to annotate large proteomic datasets and identify unsuspected roles for the ECM in pathophysiological processes. Last, to aid the dissemination and usage of the proposed definition and categorization of the *Drosophila* matrisome by the scientific community, our list has been made available through three public portals: The Matrisome Project, FlyBase, and GLAD.

## 1. Introduction

The extracellular matrix (ECM) is an assembly of hundreds of proteins that structurally supports and biochemically regulates the cells it surrounds [1,2]. The ECM organizes the tissues of all metazoans [3]. It plays a role in a number of biological processes, from development and homeostasis [4–6] to pathological processes including fibrosis and cancer [4,7,8]. With a growing interest from the scientific community in the ECM and the emergence of high-throughput technologies generating large datasets came the realization that a robust definition of the proteins contributing to the formation of the ECM was needed. We thus defined the matrisome of human and mouse [9–11]. This was achieved by developing a computational approach based on protein sequence analysis using key structural features of ECM proteins, including the presence of a signal peptide and specific protein domains found predominantly in ECM and ECM-associated proteins [9,12]. We further proposed to classify the matrisome into the core matrisome, which is the compendium of genes encoding proteins forming the structure of the ECM (collagens, glycoproteins, and proteoglycans), and the matrisome-associated ensemble comprising genes encoding accessory proteins and proteins involved in the remodeling of the ECM [9,10,13]. The adoption of these definitions by the scientific community has allowed the identification of ECM proteins previously unsuspected to play roles in physiological or pathological processes [14–16] and of ECM signatures in –omic datasets predictive, for example, of cancer patient outcomes [7,17–19]. This prompted us and others to further define the matrisome of several model organisms: zebrafish [20], *Caenorhabditis elegans* [21], and planarians [22].

In recent years, there has been a surge of interest in using the genetic tractability of *Drosophila melanogaster* to identify fundamental mechanisms underlying ECM assembly, structure, and function, since several ECM proteins and processes contributing to the formation and assembly of the ECM are conserved between *Drosophila* and other organisms [23–25]. This surge is most evident in studies of basement membrane (BM) biology [26,27]. BM is an ancient and highly conserved ECM that lines the basal surface of epithelial and endothelial tissues and surrounds muscles, adipose tissue, and nerves [26,28,29]. Studies using *Drosophila* have made particularly strong contributions to our understanding of BM secretion and assembly [30–45], and the role BMs play in shaping tissues during development [35,42,46–53]. They have also shown how BMs heal after injury [54,55] and how they regulate the immune response [56–60]. More recently, work in *Drosophila* has introduced a new role for BM proteins in intercellular adhesion [31].

Although the core BM proteins (type IV collagens, laminins, heparin sulfate proteoglycans, and nidogens) are well known, proteomic studies have revealed that BMs can harbor numerous accessory proteins that vary by tissue [14,61,62]. A comprehensive list of these proteins will provide an important tool for *Drosophila* researchers as they continue to probe the diverse roles BMs play in animal development and physiology.

*Drosophila* also have ECMs that are unique to arthropods and are therefore not found in any other organism for which the matrisome has been defined. These include: the chitin-based cuticle that forms the animal’s exoskeleton and lines the lumens of the foregut and hindgut [63–65]; non-cuticular, chitin-based ECMs that line the lumens of the trachea, salivary glands, and midgut [64,66]; the eggshell that protects the developing embryo [67,68]; and the salivary glue that is produced by the larva to affix the pupa to a surface [69]. Defining the list of proteins that comprise these *Drosophila*-specific ECMs will provide a reference dataset for the arthropod clade and aid with the annotation of large proteomic datasets, including the developmental proteome of *Drosophila* [70]. Moreover, because insects can be both disease vectors and agricultural pests, these data could provide an important source of information to combat these threats to human welfare.

Here, we define the *in-silico* matrisome for *Drosophila melanogaster*. To this end, we developed a computational pipeline that combines orthology comparison, protein sequence analysis, interrogation of experimental proteomic data, and literature search (Figure 1) and identified 641 genes that comprise the *Drosophila* matrisome. We further classified these 641 genes into different structural and functional categories based on the model we have proposed for the matrisomes of other organisms [9,20,21]. We then describe the deployment of our list and terminology in the Matrisome Project website (http://matrisome.org) and in two databases, FlyBase [71,72] and GLAD [73], broadly used by the *Drosophila* community. Last, we illustrate how this new resource can be used to annotate –omic datasets.

**Figure 1.**
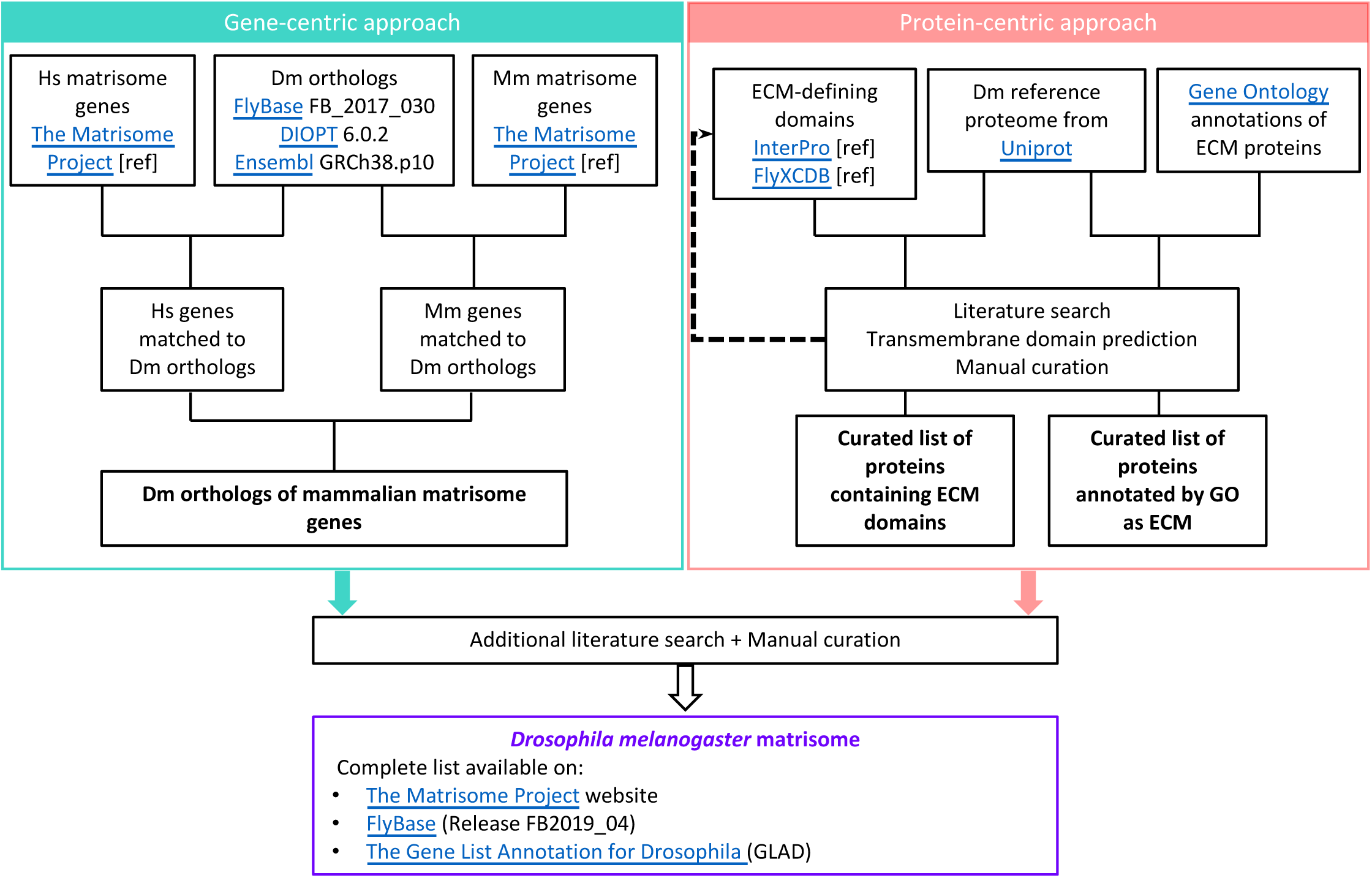
Bioinformatic workflow to define the *in-silico* matrisome of *Drosophila melanogaster*. The databases Flybase, DIOPT, and Ensembl were interrogated with the full list of human and mouse matrisome and matrisome-associated gene symbols. Selected InterPro domains, including domains characteristic of collagens, proteoglycans, ECM-affiliated proteins, and cuticle-binding proteins (see Supplementary Table S4) were used to identify ECM-domain-containing proteins in the reference proteome. The Gene Ontology annotations related to the ECM were then used to identify previously-annotated ECM components. Finally, selected published literature using proteomic and/or bioinformatic methods, as well as reviews on the subject, were searched to identify ECM proteins not identified by the orthology-based or protein-sequencing methods. These data were combined and manually curated to generate the first complete *Drosophila* matrisome.

## 2. *In-silico* definition of the *Drosophila melanogaster* matrisome

### 2.1. Identification of Drosophila orthologs of human and mouse matrisome genes

We first set out to identify the *Drosophila* orthologs of human and mouse matrisome genes. The databases Flybase (FB2017_03, released June 2017) [74], DIOPT (Version 6.0.2, released June 2017) [73], and Ensembl (Ensembl 89, released May 2017) [75] were interrogated with the full list of human and mouse core matrisome and matrisome-associated gene symbols from each of the six categories of ECM components defined by The Matrisome Project (Figure 1) [11]. The genes retrieved by each of the three databases (**Supplementary Table 1A and 1B**) were compiled to obtain a list of all predicted *Drosophila* orthologs of human and/or mouse matrisome genes (**Supplementary Table 1C**). The results of this approach led to the identification of 834 putative *Drosophila* matrisome orthologs. Of these genes, 114 were orthologous to a human gene but not a mouse gene, whereas 51 were orthologous to a mouse gene but not a human gene. There were 296 human genes with no *Drosophila* ortholog (**Supplementary Table 2A**) and 340 mouse genes with no *Drosophila* ortholog (**Supplementary Table 2B**).

### 2.2. Protein-domain-based approach to identify additional Drosophila matrisome proteins

Since it is well known that flies also have a large number of ECM proteins that do not have mammalian orthologs (*see Introduction*), we next used the UniProt *Drosophila* reference proteome (downloaded August 10, 2017) [76] to further expand our search for matrisome components (**Supplementary Table 3A)**. Taking advantage of the conserved domain-based nature of ECM proteins [12], we selected InterPro domains which were previously used to identify human and mouse matrisome proteins [9,10], including domains characteristic of collagens, proteoglycans, and ECM-affiliated proteins, to search for ECM-domain-containing proteins in the *Drosophila* proteome (**Supplementary Table 4A**). We also included in the search three domains characteristic of proteins involved in the production and maintenance of chitin-based ECMs: insect cuticle protein (IPR000618), chitin-binding domain (IPR002557), and chitin-binding type R&R consensus (IPR031311) [77]. Although three ECM domains were initially used to search the UniProt *Drosophila* reference proteome, the domain chitin-binding type R&R consensus (IPR031311) was found to be redundant with the domain insect cuticle protein (IPR000618) for the identification of the 213 *Drosophila* proteins. (**Supplementary Table 4B**). To complete the list of proteins composing the *Drosophila* cuticle we further interrogated CuticleDB, a database of structural components of arthropods identified experimentally or through protein sequence analysis [78]. This allowed us to retrieve an additional 7 genes (CG13670, CG7548, CG8541, CG8543, Cpr65Ax1, Edg91, Lcp6) that were added to the class of cuticular proteins.

Using this method, we identified 353 *Drosophila* proteins with ECM domains: 140 using domains previously used to identify mammalian matrisome proteins and an additional 213 using domains characteristic of *Drosophila* proteins (**Supplementary Table 4B**). We compared the list of proteins identified with human matrisome domains to the proteins identified via gene orthology and found that 49 of the 140 proteins discovered by domains characteristic of ECM proteins (35%) were not previously identified using the orthology approach (**Supplementary Table 4C**).

### 2.3. Gene-Ontology-based approach to identify additional Drosophila matrisome proteins

The *Drosophila* proteome retrieved from UniProt is also annotated with Gene Ontology (GO) – Cellular Component terms describing the intra- and extracellular localization of proteins [79,80]. The Gene Ontology terms extracellular matrix (GO:0031012), extracellular region (GO:0005576), extracellular space (GO:0005615), basement membrane (GO:0005604), and proteinaceous extracellular matrix (GO:0005578) were used to identify ECM components. The term proteinaceous extracellular matrix was found to be redundant with the term extracellular matrix, but the other four terms made significant contributions to the breadth of the search, which retrieved 1308 proteins from the *Drosophila* proteome (**Supplemental Table 3B**). As GO annotations have been found previously to lack specificity to define ECM components [10], analysis of all proteins identified by GO annotation was performed using the Phobius signal peptide predictor [81]. Proteins that lack a signal peptide and did not exhibit other significant ECM characteristics were excluded along with proteins predicted to be cytoplasmic, proteins with multiple transmembrane domains, and proteins with contradictory GO annotation such as cytosolic (GO:0005829) or lysosomal (GO:0005764) localization.

### 2.4. The Drosophila matrisome is composed of 641 genes

The three computational approaches described above identified 1,585 genes encoding potential *Drosophila* matrisome proteins. We then consulted selected published papers using proteomic and bioinformatic methods, read reviews on *Drosophila* ECM, and made direct queries of FlyBase [71]. This final curation step allowed us to identify some genes encoding ECM proteins that were missed by our computational screen and to eliminate some genes identified computationally for which experimental evidence does not support their classification as matrisome components (see below). The combined result of these analyses is the generation of the *Drosophila* matrisome, comprised of 641 genes (**Supplemental Table 5**). Interestingly, this number represents 4% of the 15,500 protein-coding genes in the *Drosophila* genome, which is comparable to the percentage of the genome encoding ECM proteins in humans, mice, zebrafish, and *C. elegans* [9,20,21] and is likely to be similar to the proportion of matrisome genes in the planarian genome [22,82]. Below, we describe how these 641 genes have been classified into matrisome categories based on their structure, localization, and/or function.

#### 2.4.1. Classification of Drosophila genes orthologous or homologous to mammalian matrisome genes

Genes with orthology or homology to human genes were categorized based on the previously proposed mammalian matrisome divisions (core matrisome or matrisome-associated) and categories (collagens, glycoproteins and proteoglycans for the core matrisome, and ECM-affiliated proteins, ECM regulators and secreted factors for matrisome-associated components) [9]. The *Drosophila* matrisome contains 34 core matrisome genes: 4 collagens, 27 glycoproteins, and 3 proteoglycans, the majority of which are orthologous to mammalian core matrisome genes (**Supplementary Table 5A and 5B** and Figure 2A). Only 1 collagen (pericardin) [83,84] and 6 glycoproteins (artichoke [85], anachronism [86], Defense protein l(2)34Fc, glutactin [87], tiggrin [88], and tenectin [89,90]) were *Drosophila*-specific.

**Figure 2.**
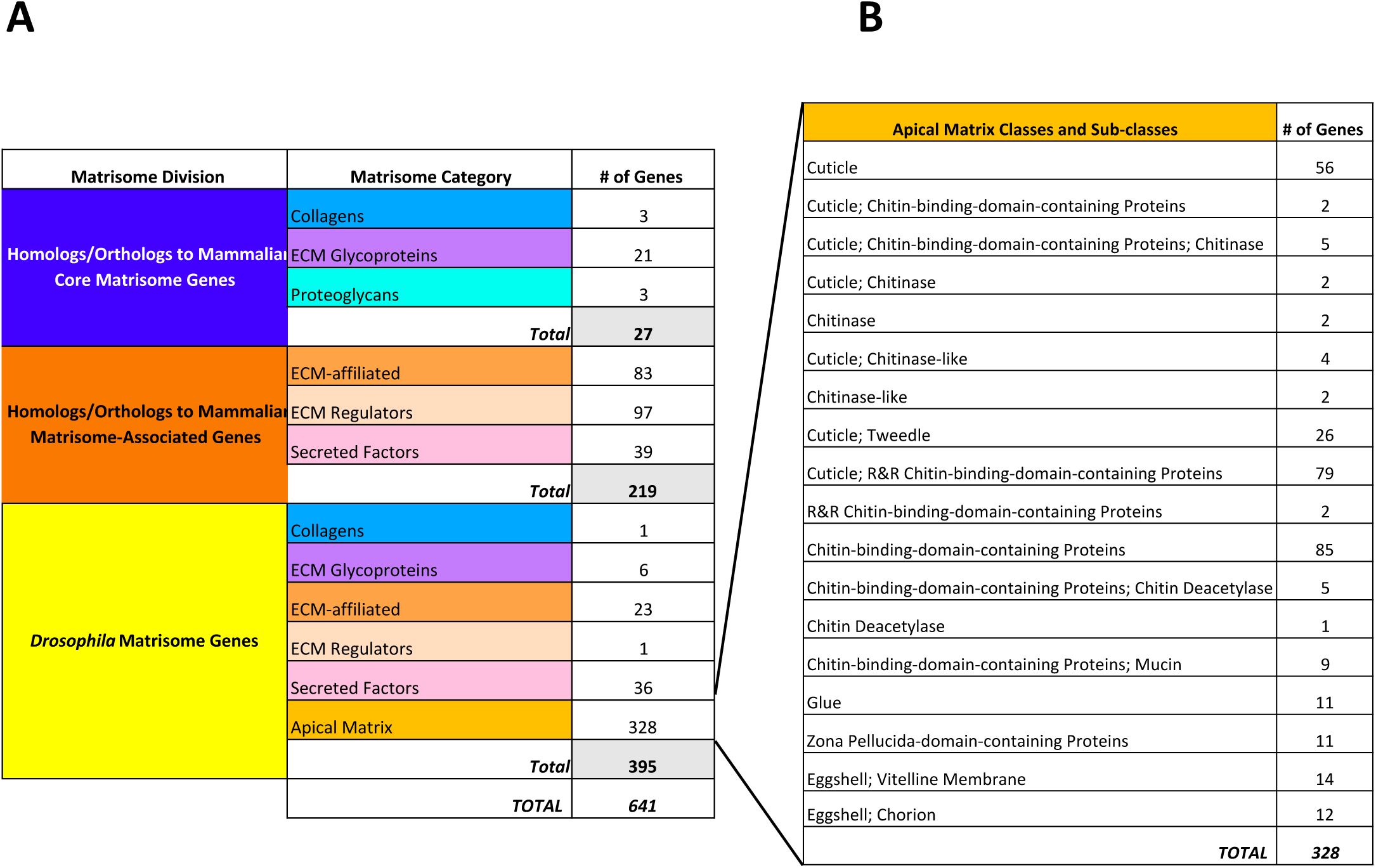
The *Drosophila* matrisome. **(A)** The *Drosophila* matrisome is made up of 641 genes. Of these, 27 are homologs/orthologs to mammalian core matrisome genes, 219 are homologs/orthologs to mammalian matrisome-associated genes, and the remaining 395 are specific to *Drosophila*. These genes are then divided into either categories which we have previously defined, or the newly proposed apical matrix category. **(B)** The genes that encode proteins that make up the apical matrix of *Drosophila* were further divided into classes and sub-classes.

In addition to core matrisome genes, we predict that the *Drosophila* genome encodes 279 matrisome-associated genes, including 219 that are orthologous or homologous to mammalian genes (**Supplementary Table 5A and 5B** and Figure 2A).

We previously defined ECM-affiliated proteins as proteins either somewhat structurally related to core ECM proteins or that have been found experimentally to be associated with the ECM in detergent-insoluble fractions of tissue lysates by proteomics [9,11]. Our computational approach predicts that 106 *Drosophila* genes encode ECM-affiliated proteins. Among these are galectins, C-type lectins (structurally characterized by three InterPro domains, IPR001304, IPR016186, and IPR016187), mucins, and semaphorins, some of which are orthologous or homologous to mammalian genes (**Supplementary Table 5A**). In addition, we classified under this category 6 collagen-triple-helix repeat-containing proteins and 9 fibrinogen-domain-containing proteins with no clear mammalian orthologs.

The ECM regulators category groups enzymes participating in the synthesis or remodeling of the ECM together with the regulators of these enzymes (including inhibitors). We identified 98 ECM regulators (**Supplementary Table 5**), including matrix metalloproteinases [25], cathepsins, ADAMs, and two orthologs of the recently identified serine/threonine kinase family Fam20 [91]. Our study also identified a total of 24 prolyl-4-hydroxylases (P4Hs). Prolyl-4-hydroxylases catalyze the formation of hydroxyprolines [24,92–94]. The most well-recognized role of this post-translational modification is to stabilize collagen triple-helical structures. Interestingly, and as previously noted [24], the human genome encodes 44 collagen genes and 3 P4Hs, whereas the *Drosophila* genome encodes only 4 collagen genes, 6 collagen-triple-helix repeat-containing proteins and yet 24 P4Hs. Both previous work [94] and interrogation of The National Human Genome Research Institute **mod**el organism **ENC**yclopedia **O**f **D**NA **E**lement (modENCODE) database [95] indicate that the P4Hs are expressed in a tissue-specific manner and at different developmental stages. Whether P4Hs have additional substrates in *Drosophila* remains to be determined.

Last, we previously included secreted factors in our definition of the matrisome, since the ECM is recognized as a reservoir of growth factors and other soluble factors [96]. These 75 proteins (**Supplementary Table 5**) were defined using a combination of orthology or homology annotations, GO terms, literature references, and the presence of characteristic domains not previously used to define secreted factors but identified from the examination of the *Drosophila melanogaster* extracellular domain database (FlyXCDB, [97]). These domains are the PDGF/VEGF domain (IPR000072), the Spaetzle domain (IPR032104), the von Willebrand factor type C (IPR029277), the insulin-like domain (IPR016179), the eclosion hormone domain (IPR006825), and the interleukin-17 family domain (IPR010345).

#### 2.4.2. Classification of Drosophila genes with no mammalian orthologs or homologs

Since the chitin-based ECM, eggshell, and salivary glue are all secreted from the apical side of epithelial tissues, we classified both the structural and regulatory proteins associated with these ECMs under a category termed “Apical ECM” (**Supplementary Table 5** and Figure 2B). As a group, these proteins comprise nearly 50% of the *Drosophila* matrisome. To further subdivide this diverse group of proteins, an additional level of classification was created to reflect their respective proteins’ domain structure, enzymatic function, or localization (**Supplementary Table 5, column C**). The chitin-binding-domain-containing proteins and R&R chitin-binding-domain-containing proteins families refer to proteins containing InterPro domains IPR002557 and IPR031311, respectively. The Chitinase and Chitinase-like families also have a group of defining domains: the chitinase II domain (IPR011583) and three glycoside hydrolase domains (IPR029070, IPR001223, and IPR017853). The Tweedle family represents the only proteins with the domain DUF243 (IPR004145) [98]. Chitin deacetylases were identified based on the presence of a glycoside hydrolase/deacetylase domain (IPR011330). A group of 11 zona-pellucida-domain-containing proteins was also identified [99]. These proteins have a shared structural attribute, the zona pellucida domain (IPR001507), which we originally used to identify core components of the mammalian matrisome. However, since zona-pellucida-domain-containing proteins do not present clear orthology or homology with mammalian proteins, we classified them apart.

Groups without clear structural similarities were classified by other means. Proteins of the cuticle that did not meet the definitions above were classified by their shared GO term, chitin-based cuticle development (GO:0040003). Included in this class were also a number of genes reported to be cuticle proteins of low complexity [100,101]. The eggshell superfamily includes two protein classes, corresponding to the vitelline membrane and chorion layers of this ECM, respectively [68,102,103]. The vitelline membrane proteins were defined by GO term or literature search. The chorion proteins had all previously been assigned chorion-related GO terms and chorion-related protein names, except Cp38 which has chorion in the name and is cited [102]. Finally, 11 proteins including new-glue and salivary glue secretion proteins, Eig71Ee [69,104], and the newly identified tandem paralog of Sgs5 (FBgn0038523) [105] were classified as glue proteins.

## 3. Accessing the *Drosophila* matrisome and utilizing it to annotate large datasets

### 3.1. The Drosophila matrisome is available from three sources

To facilitate the use of our definition and categorization of the *Drosophila* matrisome by the scientific community, the list devised here has been made available through three public platforms. Similar to the matrisome lists of human, mouse, zebrafish and *C. elegans*, the *Drosophila* matrisome list can be found on the Matrisome Project website (http://matrisome.org [11]. Moreover, it has been implemented in two databases widely used by the *Drosophila* community. The *Drosophila* matrisome is be available within the “Gene Groups” section of the August 2019 release of FlyBase (FB2019_04), which is the most comprehensive source of genetic information for this model organism [71,72,106]. In addition, as a result of the Matrisome analysis, two new terms were added to the Gene Ontology Cellular Component aspect: chitin-based extracellular matrix (GO:0062129) and adhesive extracellular matrix (GO:0062130), allowing more precise GO annotation of the constituents of these specific types of ECM. All *Drosophila* cuticle proteins and glue genes have now been annotated with these respective terms in FlyBase.

The *Drosophila* matrisome is also available in the Gene List Annotation for *Drosophila* (GLAD) database, which is maintained by the Perrimon laboratory to enhance the utility of the cell-based RNAi screening (DRSC) and *in vivo* fly RNAi (TRiP) collections for the community [73]. For consistency with the current GLAD nomenclature, the matrisome forms a new gene list/group; the matrisome divisions are listed as sub-groups, the categories as sub-sub-groups, and the families are listed under comments.

### 3.2 The Drosophila matrisome provides a powerful tool to annotate large datasets

One powerful application of the matrisome list for any species is in the annotation of large-omic datasets [11]. Thus, as a proof of principle, we used the newly defined *Drosophila* matrisome to re-evaluate two recently published datasets that focus heavily on ECM-associated proteins. In the first study, Baycin-Hizal and colleagues identified 399 N-glycosylated proteins of the *Drosophila* head region using solid phase extraction of N-linked glycopeptides coupled to LC-MS/MS [107]. They reported that 4.5% of the proteins identified experimentally in their study were part of the ECM. We found, however, that 13% of the proteins they identified (which included 8 of the 26 glycoproteins and 2 of the 3 proteoglycans we have predicted) are in fact matrisome proteins, more than double the original number. In the second study, Sessions and colleagues reported changes in the abundance of ECM proteins in the *Drosophila* heart during aging [108]. 104 of the proteins detected were identified as ECM proteins using the Software Tool for Rapid Annotation and Differential Comparison of Protein Post-Translational Modifications (STRAP PTM) developed by Spender and colleagues. Of these 104 proteins, 27 are part of the matrisome, whereas 77 are not. Examination of these 77 proteins revealed that most are in fact localized intracellularly, with little evidence to support that they are ECM components. We retrieved the raw mass spectrometry data from the ProteomeXchange repository (PXD006120) and reannotated the data using the matrisome list. We identified a total of 46 matrisome proteins, finding 19 additional proteins not originally annotated as belonging to the ECM. Together, these two examples demonstrate the power of our matrisome list to comprehensively annotate large experimental datasets. We thus propose that the use of our annotations and nomenclature would assist in the comprehensive identification of ECM signatures contributing to cellular, physiological and pathological phenotypes.

## 4. Conclusion

We defined here the *in-silico Drosophila melanogaster* matrisome. In addition to reporting the identification of 641 genes encoding ECM and ECM-associated proteins, we further propose their comprehensive classification according to structural and/or functional features. We hope that this list and nomenclature will aid with the annotations of large datasets, and thus further our understanding of the roles of the ECM in fundamental biological processes and pathophysiology.

## Acknowledgements

We would like to thank the members of the Naba Lab for critical reading of the manuscript. We would also like to thank our long-time collaborator Karl Clauser for his help analyzing the raw proteomic data from the Sessions et al., 2016 study.

Last, we would like to thank Dr. Helen Attrill (University of Cambridge), curator of FlyBase, for her help with the curation of the matrisome list and for its implementation to FlyBase, and Dr. Claire Yanhui Hu from the *Drosophila* RNAi Screening Center (DRSC) and Transgenic RNAi Project (TRiP) in Norbert Perrimon’s laboratory (Harvard Medical School) for implementing our list to GLAD.

## SUPPLEMENTARY TABLE LEGENDS

**Supplementary Table 1.** *Drosophila* orthologs to human and mouse matrisome genes

**A. and B.** List of *Drosophila* genes identified by orthology to (A) human and (B) mouse matrisome genes (column A) and the matrisome category of their human or mouse orthologs (columns B-G). 783 orthologs to human matrisome genes and 720 orthologs to mouse genes were discovered.

**C.** List of all *Drosophila* orthologs to human or mouse matrisome genes (column A). Columns B and C indicate whether they were orthologous to human genes, mouse genes, or both. A total of 834 *Drosophila* orthologs to human and mouse matrisome genes were discovered.

**Supplementary Table 2.** Human and mouse genes with no *Drosophila* ortholog

**A.** Human genes with no *Drosophila* orthologs predicted.

**B.** Mouse genes with no *Drosophila* orthologs predicted.

**Supplementary Table 3.** UniProt reference proteome

**A.** The UniProt *Drosophila* reference proteome was retrieved August 10th, 2017. To allow for cross-referencing with our gene-specific list of orthologs each gene in the reference proteome was assigned a primary gene name (column E) based on the first gene name listen under column F.

**B.** The reference proteome was interrogated with three Gene Ontology terms, extracellular matrix (GO:0031012), extracellular region (GO:0005576), and basement membrane (GO:0005604), to obtain a set of 1308 putative matrisome proteins.

**Supplementary Table 4.** InterPro domains

**A.** List of ECM domains (column C) used to detect *Drosophila* matrisome proteins and their matrisome categories (columns A and B).

**B.** All *Drosophila* proteins retrieved from the UniProt *Drosophila* reference proteome and the InterPro domain families (column A) used to retrieve them.

**C.** Comparing the proteins identified by the protein-domain-based approach to the list of proteins previously generated by the orthology-based approach demonstrated that 49 novel proteins were identified.

**Supplementary Table 5.** The complete annotated *Drosophila melanogaster* matrisome

**A.** Drosophila matrisome genes are organized by matrisome division, category, and class (columns A-C). Each gene is associated with a unique FlyBase gene ID (columns D and E). Also provided are all UniProt IDs associated with the gene and a protein name pulled from the UniProt *Drosophila* reference proteome (columns F-H). InterPro domains, Gene Ontology terms, and alternative gene names were also obtained from InterPro (columns I-K, M). Genes added to the matrisome via the gene-centric, orthology-based approach have their human orthologs or homologs listed (column L).

**B.** The number of genes in each matrisome division and category. All combinations of classes of the apical matrix category of genes are also shown, as well as the same genes divided into their most granular (listed first in A, column C) classes or families (see also Figure 2).

